# Host adaptation through hybridization: Genome analysis of triticale powdery mildew reveals unique combination of lineage-specific effectors

**DOI:** 10.1101/2021.05.06.442769

**Authors:** Marion C. Müller, Lukas Kunz, Johannes Graf, Seraina Schudel, Beat Keller

## Abstract

The emergence of new fungal pathogens through hybridization represents a serious challenge for agriculture. Hybridization between the wheat mildew (*Blumeria graminis f.sp. tritici*) and rye mildew (*B.g. f.sp. secalis*) pathogens have led to the emergence of a new mildew form (*B.g. f.sp. triticale*) growing on triticale, a man-made amphiploid crop derived from crossing rye and wheat which was originally resistant to the powdery mildew disease. The identification of the genetic basis of host-adaptation in triticale mildew has been hampered by the lack of a reference genome. Here we report the 141.4 Mb reference assembly of *B.g. triticale* isolate THUN-12 derived from long-read sequencing and genetic map-based scaffolding. All eleven *B.g. triticale* chromosomes were assembled from telomere-to-telomere and revealed that 19.7% of the hybrid genome was inherited from the rye mildew parental lineage. We identified lineage-specific regions in the hybrid, inherited from the rye or wheat mildew parental lineages, that harbour numerous bona fide candidate effectors. We propose that the combination of lineage-specific effectors in the hybrid genome is crucial for host-adaptation, allowing the fungus to simultaneously circumvent the immune systems contributed by wheat and rye in the triticale crop. In line with this we demonstrate the functional transfer of the *SvrPm3* effector from wheat to triticale mildew, a virulence effector that specifically suppresses resistance of the wheat *Pm3* allelic series. This transfer is the likely underlying cause for the observed poor effectiveness of several *Pm3* alleles against triticale mildew and exemplifies the negative implications of pathogen hybridizations on resistance breeding.

## Main Text

The emergence of new fungal pathogens on crops poses a serious problem for agriculture. One mechanism by which fungal pathogens can adapt to a new host is hybridization, by which two different pathogen lineages recombine their gene complement enabling them to infect a new host species (Thines, 2019). Despite several reports of hybridizations in plant pathogens, host adaptation by hybridization is poorly understood and the genetic loci involved remain unidentified (reviewed in (Stukenbrock, 2016).

A prominent example of a host-adaptation through hybridization is the recent emergence of the triticale powdery mildew (*Blumeria graminis f,sp, triticale*) on the cereal crop triticale in the year 2001 (Walker *et al*., 2011). *B.g. triticale* was found to be the result of hybridization events between the highly host-specific fungal sub-lineages of wheat *(B.g. tritici*) and rye (*B.g. secalis*) powdery mildew (Menardo *et al*., 2016). The hybridization on the pathogen side mirrored the breeding history of its new host, which is an amphiploid resulting from a cross of the two cereal crops wheat (*Triticum. sp*.) and rye (*Secale cereale*) (Oettler, 2005). Triticale has seen an increase in cultivation in the past decades especially in Europe (Arseniuk, 2014). Unfortunately, disease outbreaks of powdery mildew on triticale have increased both in number and severity and corroborates the need for resistance breeding in this crop (Mascher *et al*., 2006; Kowalczyk *et al*., 2011; Arseniuk, 2014).

Previous analyses of the triticale powdery mildew hybrid pathogen were based on the wheat mildew reference genome (Menardo *et al*., 2016; Praz *et al*., 2018). This approach suffers from its bias to the gene content of one of the parental lineages of the hybrid. This is particularly relevant since effectors, small, secreted proteins that are encoded by hundreds of genes in powdery mildew genomes, reside in highly dynamic gene clusters that often show copy number variation or lineage specific expansions (Menardo *et al*., 2017; Frantzeskakis *et al*., 2018; Müller *et al*., 2019). Due to their dual role in infection as putative virulence factors suppressing the host defences as well as avirulence factors recognised by major resistance genes, effector genes represent prime candidates for host-specificity factors (Li *et al*., 2020). In this study we present a chromosome-scale assembly of triticale powdery mildew that will lay the foundation to understand host adaptation through hybridization in the cereal powdery mildew pathosystem.

We sequenced *B.g. triticale* isolate THUN-12 using PacBio technology to a sequencing depth of 180X to establish a reference assembly of the hybrid powdery mildew. The resulting assembly was 141.1 Mb in size and consisted of 51 contigs. We used the previously described scaffolding method based on the genetic map of the fully sequenced mapping population of THUN-12 X *B.g. tritici* Bgt_96224 (Müller *et al*., 2019) to scaffold the contigs into eleven chromosomes (Table S1). Three of the eleven chromosomes, namely Chr-03, Chr-06 and Chr-10, were assembled in a single contig in the THUN-12 assembly. The remaining chromosomes were scaffolded with a maximum of three scaffold gaps. In addition, the genetic map allowed to correct three assembly errors, in which chromosomes arms originating from different chromosomes were fused in the centromeric regions. With a transposable element content of 69.4%, the *B.g. triticale* genome exhibits a characteristic genome organization with high repeat content, reminiscent of other sequenced *B. graminis* isolates. (Frantzeskakis *et al*., 2018; Müller *et al*., 2019) We found a single region per chromosome with a distinct transposable element content consisting of non-long-terminal-repeat retrotransposons, corresponding to the genetic centromere (Figure S1). In addition, we found stretches of telomeric repeats TAACCC at all ends of the eleven chromosomes (Figure S1). The presence of telomeres and the centromere on each chromosome together with the low number of scaffold gaps indicates high completeness of all eleven chromosomes resolved in the *B.g. triticale* assembly. This represents a significant improvement to previous high-quality assemblies of the *Blumeria graminis* species, which still contain hundreds of gaps or unresolved chromosomal regions (Frantzeskakis *et al*., 2018; Müller *et al*., 2019).

The high-quality genome of THUN-12 allowed us to study the signature of hybridization between *B.g. tritici* and *B.g. secalis* at the chromosomal level. As described in (Menardo *et al*., 2016) we used fixed polymorphisms between *B.g. secalis* and *B.g. tritici* to attribute genomic segments in THUN-12 to the two parental sub-lineages ((Menardo *et al*., 2016). We found that 80.3% of the THUN-12 genome is inherited from wheat mildew and 19.7% was inherited from rye mildew (Figure 1, Table S1). The observed higher percentage of segments with wheat mildew origin in the THUN-12 isolate is consistent with the previously proposed scenario in which the first rye/wheat mildew hybrids backcrossed to wheat mildew after the initial hybridization (Menardo *et al*., 2016). During meiosis, at least one crossover takes place between homologous chromosomes (Marston & Amon, 2004). Indeed, we found at least one recombination event between wheat and rye mildew per chromosome, suggesting efficient chromosomal pairing in all chromosomes between the two parental lineages during the initial hybridization. Strikingly, the size of segments inherited form *B.g. secalis* varied considerably and proportional contribution to chromosomes ranged from 6.0% on Chr-09 to a maximum of 40.9% on Chr-04 (Figure 1, Table S1). In addition, the location of the *B.g. secalis* segments is highly variable between chromosomes, for instance, eight of the eleven chromosomes inherited at least one telomeric sequence form *B.g secalis*. In contrast, Chr-07 contains a single, larger segment inherited from *B.g. secalis* that overlaps with the genetic centromere. The centromeres of all other chromosomes were inherited from *B.g. tritici*. To what degree the differential proportion of *B.g. secalis* segments per chromosome are the result of differential contribution to host-adaptation or purely stochastic based on the limited amount of recombination between the parental strains remains to be determined in further studies.

**Fig. 1.**
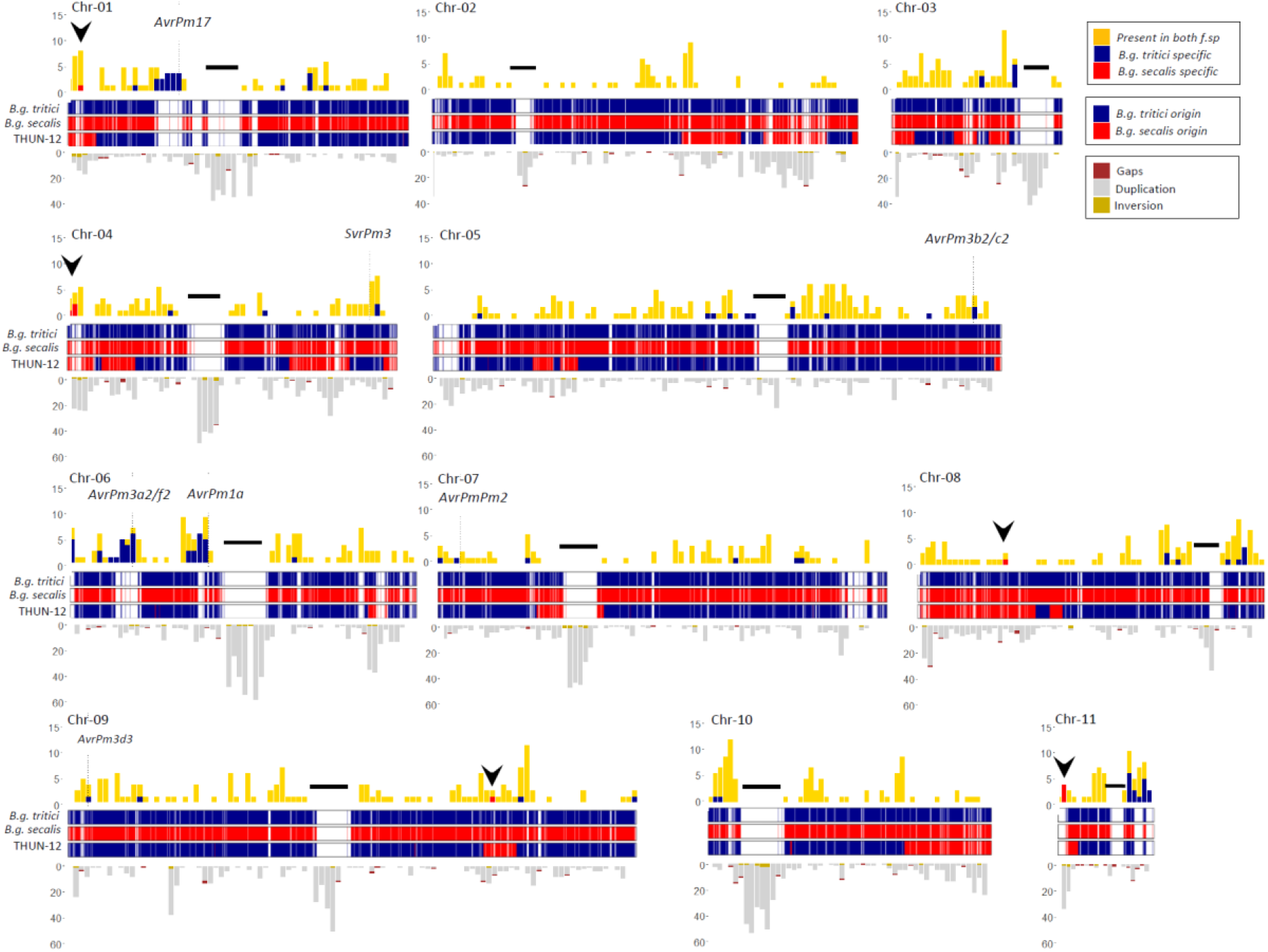
Overview of the eleven chromosomes of *B.g. triticale* THUN-12. The first track shows the distribution of candidate effector genes along the eleven chromosomes. Candidate effectors present in both parental species *f.sp. tritici* and *f.sp. secalis* are indicated in yellow; candidate, lineage-specific effector genes of *B.g. secalis* isolates are indicated in red and candidate, lineage-specific effector genes of *B.g. tritici* are indicated in blue. Black arrowheads indicate the position of regions depicted in Figure 2. The second track indicates the genomic origin of the THUN-12 sequence, based on fixed polymorphisms between *B.g. secalis* and *B.g. tritici*. SNPs originating from *B.g. secalis* are indicated in red, SNPs originating from *B.g. tritici* are represented in blue. For each chromosome the position of the genetic centromere is indicated by a black bar. The third track indicates number of putative alignment breaks between *B.g. triticale* THUN-12 and *B.g. tritici* Bgt_96224 based on whole genome alignment of the chromosome-scale assemblies of the two *f.sp*. by the MUMmer software. Putative rearrangements were filtered and only rearrangements bigger than 1kb and smaller than 10kb are depicted. The alignment breaks were depicted according to the prediction of the MUMmer software as follows: brown indicates putative gaps, grey putative duplications and yellow indicates putative inversions.

The availability of the hybrid THUN-12 genome and the previously published chromosome-scale assembly of the wheat mildew Bgt_96224 (Müller et al., 2019) allowed the comparison of the hybrid genome to the wheat mildew parental lineage.. Upon whole-genome alignment of the two genomes (Figure S3) using the MUMmer software (Kurtz *et al*., 2004) 99.26% of the THUN-12 genome was aligned to the Bgt_96224 genome. We observed a high co-linearity across the eleven chromosomes in *B.g tritici* inherited regions as well as in the 27.7 Mb of sequence originating from *B. g. secalis* (Figure S3). The analysis identified only a few (<20) regions with major rearrangements, most of which overlap with the genetic centromeres and likely represent assembly artefacts (Figure S3). The centromeres consist exclusively of repetitive elements and are therefore often not completely resolved despite the long-read sequencing technology (Müller et al. 2019). In addition, we identified a large inversion on Chr-02. The inverted region in Bgt_96224 is however flanked by sequence gaps and could therefore be explained by a scaffolding error in the Bgt_96224 assembly. The most interesting major difference between the genomes is a telomeric region of Chr-11 which is absent in the assembly of Bgt_96224 (Figure S2) and therefore represents a candidate lineage-specific region in THUN-12 which was inherited from *B.g. secalis* (see below). When we considered smaller rearrangements (>1kb, <10kb) predicted by the MUMmer software, we found evidence for alignment breaks that constitute inversions, gaps and duplications on all chromosome (Table S4). Segments inherited from *B.g. secalis* were significantly enriched for such rearrangements compared to segments inherited from *B.g. tritici*, which is consistent with the divergence of the two fungal lineages ca. 160’000-250’000 ya (Table S4, (Menardo *et al*., 2016). Due to the predominance of small-scale rearrangements identified by comparison to *B.g. tritici*, we concluded that the two parental f.sp. of the hybrid have a very similar overall genome structure. This is consistent with our findings in the experimental population Bgt_96224 X THUN-12 for which we found no impairment of recombination in segments inherited from *B. g. secalis*. (Müller et al. 2019). We therefore propose that the high similarity of the two genomes provided the basis for the successful hybridization of *B.g. secalis* and *B.g. tritici* that gave rise to the first *B.g. triticale* hybrids and that the occurrence of hybridization of these lineages was mainly limited by suitable host plants in the past.

The process of hybridization allows the combination of genes or genomic regions that are normally present only in one particular fungal lineage, but absent in the other. In numerous fungal plant pathogens host-adaptation has been attributed to the occurrence of lineage-specific effector proteins acting as virulence factors (i.e. allowing growth on the particular host) or avirulence factors (i.e. preventing growth due to recognition by host specific immune receptors). To identify putative effector genes in the *B.g. triticale* hybrid, we performed a de-novo gene annotation based on the previously published proteins of *B.g. tritici* and *B.g. hordei*. This resulted in the annotation of 7,993 genes including 1,011 candidate effectors that were subsequently assigned to the previously described *Blumeria* candidate effector families (Müller *et al*., 2019). To test for the parental contribution to the gene content of the hybrid we determined for each gene whether it is encoded in a segment derived from wheat or rye mildew. We found that a total of 20 % of all genes in THUN-12 were encoded within a segment of *B.g. secalis*. To define lineage-specific effectors we used genomic coverage based on resequencing data from *B.g. tritici* or *B.g. secalis* isolate to predict the absence of the THUN-12 candidate effectors in all isolates within one parental forma specialis (Note S2). We identified five loci containing candidate, lineage-specific effector genes inherited from *B.g secalis* in the telomeric regions of Chr-01, Chr-04, and Chr-11 and on the chromosome arms of Chr-08 and Chr-09 (Figure 1). Upon comparison with the Bgt_96224 genome, we could confirm the absence of these effector genes within the wheat mildew lineage (Figure 2, Note S3,). For instance, the *B.g. secalis* lineage-specific region on Chr-01 in THUN-12 contains a three-fold tandem duplication of two effectors belonging to family E135 and E001, respectively, whereas the corresponding region in Bgt_96224 contains only one gene each (Figures 2A, S6B). On Chr-04 we found a cluster of the E029 effector family with 9 members in THUN-12 and 6 members in Bgt_96224. Therefore, this cluster predates the split between *B.g. secalis* and *B.g. tritici*, but rearrangements within the cluster have led to three lineage-specific effectors in *B.g. secalis* (Figure 2B). The third example represents the above-mentioned lineage-specific region on THUN-12 Chr-11 which was found to be partially absent from the Bgt_96224 assembly. The region contains six candidate effector genes of which three belong to family E003 and are lineage-specific for *B.g secalis* (Figure 2C). The remaining two regions on Chr-08 and Chr-09 contained a single effector gene present in THUN-12 but absent from Bgt_96224 belonging to E004 and E001, respectively (Figure 2D,E). It is worth noting that several of the identified *B.g. secalis* lineage-specific candidates exhibit similarities with validated virulence or avirulence factors in *B.g. tritici* and *B.g. hordei* (Note S4). For example, several candidates belong to a group of small, highly expressed candidate effector families (termed group 1 effectors) harboring most of the virulence/avirulence factors in the *Blumeria* genus identified to date (Müller *et al*., 2019). Similarly, numerous genes are predicted to exhibit structural similarities (RNAse-like fold) to cloned avirulence genes and virulence factors (Figure S5) (Bauer *et al*., 2021). Most importantly, several lineage-specific candidates exhibit very high expression levels during the crucial phase of haustoria establishment (Figure S5), a hallmark of avirulence factors in wheat mildew (Bourras *et al*., 2019).

**Fig. 2.**
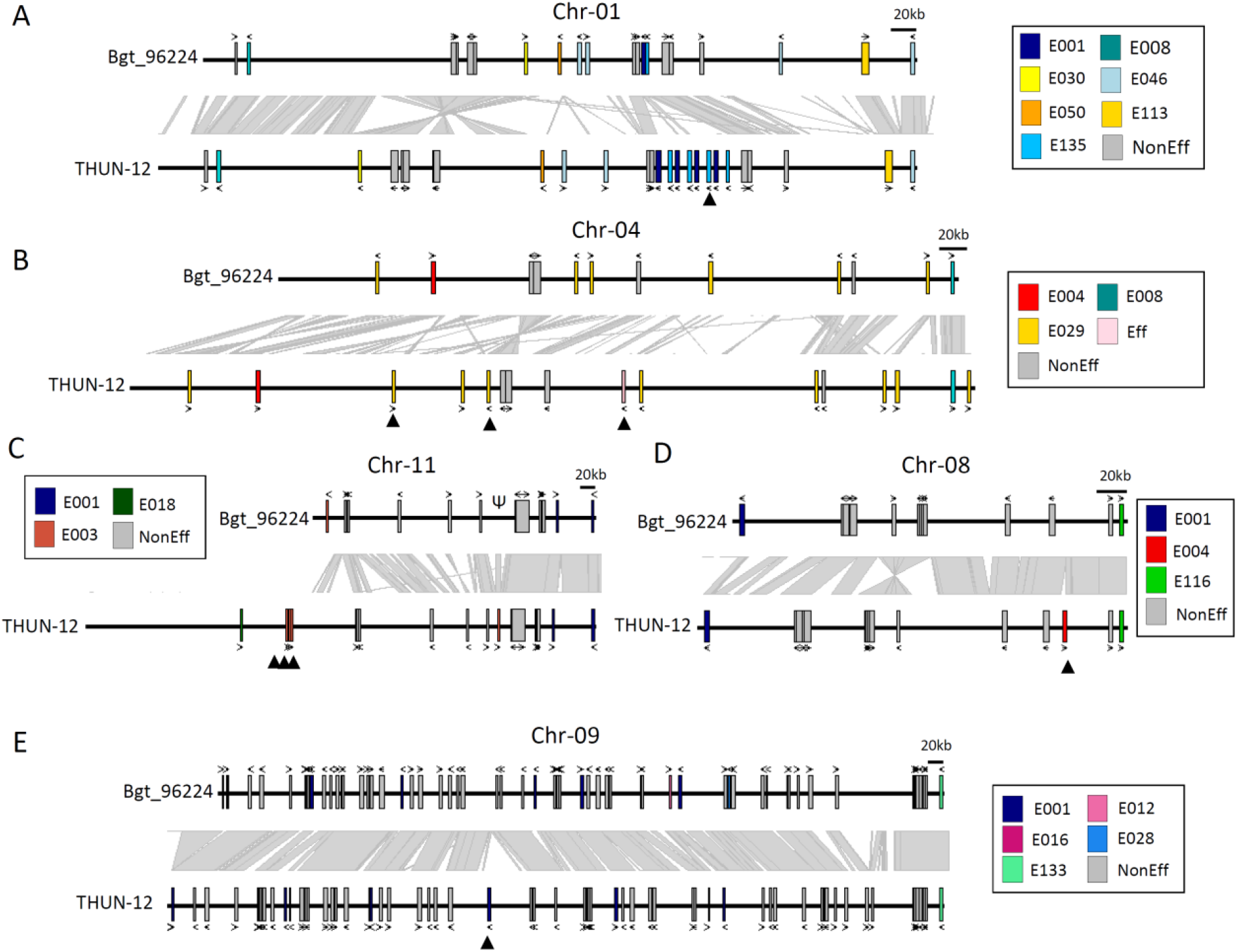
Identification of lineage-specific candidate effector genes in THUN-12 inherited from the *B.g. secalis* parental isolate. **A-E)** Candidate effectors genes are indicated by colored boxes according to their candidate effector family (Müller et al. 2019). Non-effector genes representing single copy orthologs determined by the OrthoFinder software are indicated by grey boxes. In panel **B)**, Eff indicates an effector gene not assigned to a candidate effector family. Gene direction is indicated above or below the gene. Co-linearity as determined by the MUMmer software is indicated by grey rectangles between the gene tracks. Genes for which a lineage-specific signal was detected are marked by a black arrowhead. **A-C)** represent the telomeric ends of Chr-01 **A)**, Chr-04 **B)** and Chr-11 **C)**, whereas **D-E)** represent genomic loci within the chromosome arms of Chr-08 **D)** and Chr-09 **E)**. The full description on the identification of the lineage-specific candidate effector is available in Note S3.

Using the same coverage-based approach we also identified several regions containing candidate effector genes absent from the *B.g. secalis* lineage. We found four loci with strong signatures of *B.g. tritici* lineage-specificity on Chr-01, Chr-06 and Chr-11 (Figure 1). Strikingly, three loci overlapped with the known *B.g. tritici* gene clusters harbouring *AvrPm17* (Chr-01), *AvrPm3*^*a2/f2*^ (Chr-06) and *AvrPm1a* (Chr-06) (Note S5). It was previously hypothesized that the strong selection pressure exerted by corresponding resistance genes in wheat has led to effector cluster expansions in these regions in the *B.g. tritici* lineage (Müller *et al*., 2019; Hewitt *et al*., 2020; Müller *et al*., 2021). The fourth region showing a high density of *B.g. tritici* specific effectors on Chr-11 of THUN-12 might consequently harbour a so far unidentified avirulence gene. Interestingly the *B.g. tritici* lineage-specific effectors on Chr-11 belong to the E003 family. Gene members of the same family are also present in the *B.g. tritici* specific locus on Chr-01 (*AvrPm17*, (Müller *et al*., 2021)) and the *B.g. secalis* specific region on the other arm of Chr-11 (see above). The E003 family members therefore represent prime candidates for effectors involved in host-adaptation on triticale. In summary we found that the *B.g. triticale* hybrid contains lineage-specific effectors inherited from both parental lineages. We hypothesize that these regions play a role in host-adaptation and virulence of triticale powdery mildew on the new host triticale, enabling the fungus to cope with both the rye and wheat immune system, simultaneously present in triticale.

As described above, three loci with strong evidence for a lineage-specific expansion in THUN-12 correspond to the previously identified *AvrPm1a, AvrPm17* and *AvrPm3*^*a2/f2*^ loci in wheat mildew. In addition, the regions corresponding to the other cloned wheat mildew avirulence genes, namely *AvrPm2, AvrPm3*^*b2/c*,^ *AvrPm3*^*d*3^ are also encoded in segments from wheat mildew in THUN-12. Therefore, the genomes of THUN-12 and Bgt_96224 provided us with a unique opportunity to compare all the previously described wheat mildew avirulence loci in two high quality genomes (Figure 3A-F). This comparison allowed us to confirm previous observations based on short read-sequencing data. For instance, THUN-12 contains three *AvrPm3*^*a2/f2*^ genes, as we have previously predicted for many *B.g. tritici* isolates based on genomic coverage (Müller *et al*., 2019). Furthermore, the *AvrPm3*^*b2/c2*^-locus contains an additional candidate effector in THUN-12 that is pseudogenized in Bgt_96224 by a transposable element insertion (Bourras *et al*., 2019), whereas the *AvrPm3*^*d3*-^locus exhibits several small scale insertions surrounding the avirulence gene. Also, THUN-12 contains the known deletion covering the *AvrPm2* gene, a deletion that is adaptive since it allows mildew to overcome *Pm2* mediated resistance in wheat (Praz *et al*., 2017). In addition, we could detect new variation previously undetected such as a small inversion affecting an effector in the *AvrPm1a* locus as well as a non-allelic gene conversion event in the newly identified *AvrPm17* paralogous effector pair (see (Müller *et al*., 2021). Strikingly, despite the dynamic nature of these avirulence loci, the THUN-12 isolate, with the exception of the deleted *AvrPm2*, encodes for the avirulent allele of all five avirulence genes. It was previously proposed that avirulence genes in powdery mildew exert an important virulence function on wheat (McNally *et al*., 2018; Bourras *et al*., 2019). We therefore hypothesize that the functional conservation of *B.g. tritici* avirulence effectors in *B.g. triticale* provides the hybrid pathogen with increased virulence capacity, at least in the absence of corresponding resistance genes in triticale.

**Fig. 3.**
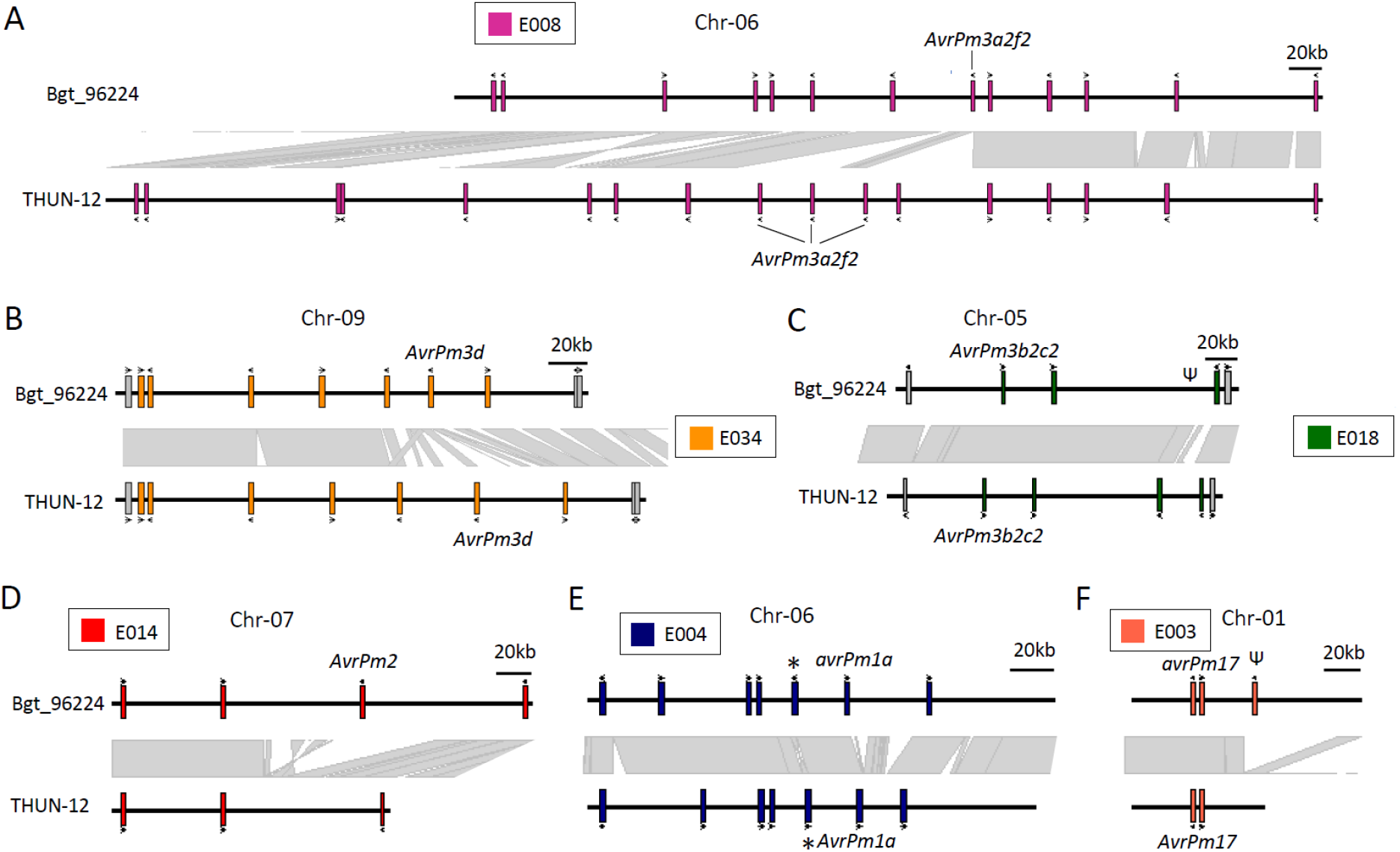
Comparison of six avirulence loci identified in wheat mildew between the *B.g. triticale* isolate THUN-12 and *B.g tritici* isolate Bgt_96224. **A-F)** The hybrid mildew THUN-12 inherited all these loci from the *B.g. tritici* parental lineage. Genes are indicated by a box, gene orientation is indicated above or beneath the box. Genome aligned as determined by the MUMmer software is indicated by grey rectangles between the gene tracks. Effector genes are indicated by colored boxes according to their candidate effector family (Müller et al. 2019). Grey boxes indicate non-effector genes. Loci were manually checked and low-confidence genes (i.e. pseudogenes, transposable elements) were removed. Asteriks in panel **E)** indicate small inversions affecting a candidate effector gene.

To test the functionality of the *B.g. tritici* avirulence genes in the hybrid genetic background, we analysed their expression during early infection on triticale at the timepoint of haustorium formation (i.e. 48hpi). Similar to the situation in wheat mildew, the avirulence genes are consistently among the highest expressed genes in THUN-12 (Figure 4A, Müller *et al*., 2021). In line with this, THUN-12 is avirulent on wheat lines containing *Pm1a, Pm17, Pm3a, Pm3b* and *Pm3d* (Figure 4B). We however observed virulent phenotypes of THUN-12 on near isogenic lines (NILs) containing *Pm3f* (weak allele of *Pm3a*) and *Pm3c* (weak allele of *Pm3b*), despite the presence of *AvrPm3*^*a2/f2*^ and *AvrPm3*^*b2/c2*^ (Figure 4B). To resolve this phenotypic/genotypic discrepancy, we performed QTL mapping based on 55 progeny of the Bgt_96224 X THUN-12 population on the *Pm3c*-containing wheat line Sonora/8*CC. We found a single QTL (LOD=9.44) on Chr-04 (Figure 4C) that corresponds to the previously described *SvrPm3*-locus in wheat mildew (Bourras *et al*., 2015; Parlange *et al*., 2015). *SvrPm3* encodes an effector that suppresses resistance mediated by *Pm3a - f*. Indeed, THUN-12 contains the active *SvrPm3* haplovariant, again encoded by a wheat mildew segment (Figures 1,4D, S6), whereas Bgt_96224 encodes the inactive allele (Bourras *et al*., 2015; Parlange *et al*., 2015). Thus, the virulent phenotype of THUN-12 on the weak *Pm3c* and *Pm3f* alleles (Brunner *et al*., 2010) can be explained by the presence of the *SvrPm3* suppressor gene, efficiently masking recognition of *AvrPm3*^*a2/f2*^ and *AvrPm3*^*b2/c2*^, thereby demonstrating the functionality of a major wheat mildew virulence factor in the hybrid genetic background. Since *B.g. secalis* encodes for an inactive *SvrPm3* haplovariant (Bourras *et al*., 2019), the active variant could only be inherited from the *B.g. tritici*. The presence of the active *SvrPm3* in the triticale mildew population likely has implications for triticale breeding. It was previously demonstrated that *SvrPm3*-based suppression is quantitative and that isolates expressing the *SvrPm3* to very high levels are able suppress also the stronger *Pm3a* and *Pm3b* alleles (Bourras *et al*., 2015; Bourras *et al*., 2019). Therefore, the presence of *SvrPm3* in the *B.g. triticale* population potentially renders the entire *Pm3*-allelic series ineffective for controlling mildew on triticale. Indeed, previous studies have found virulence proportion of over 40% towards several *Pm3*-alleles in the triticale powdery mildew population of Poland, France and Belgium (Czembor *et al*., 2013; Troch *et al*., 2013; Czembor *et al*., 2014). We propose that these results can be explained by the presence of the active *SvrPm3* in these populations.

**Fig. 4.**
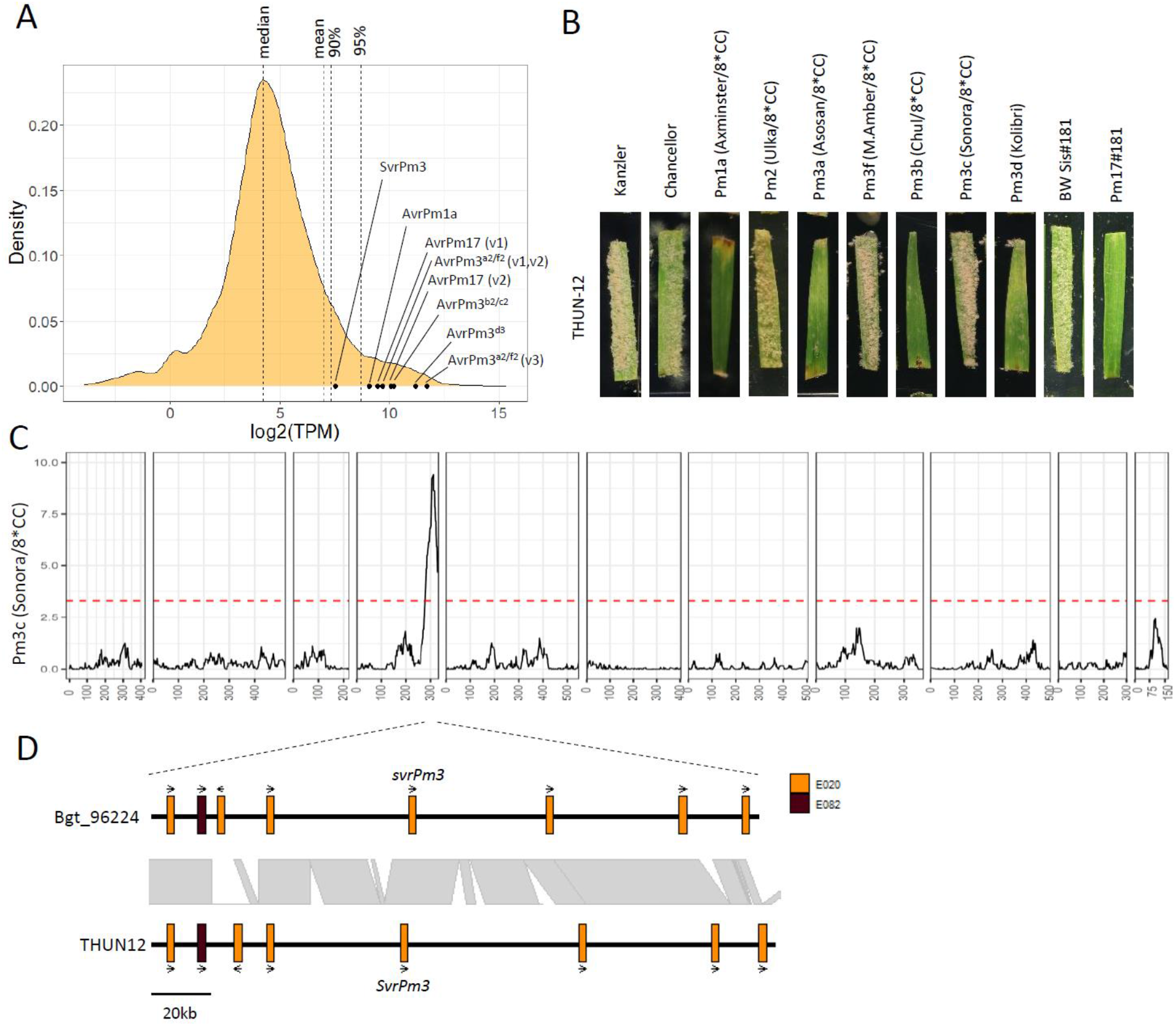
Analysis of avirulence genes in *B.g. triticale* THUN-12. **A)** Summary of gene expression of all genes in isolate THUN-12 at two days post infection (2dpi) on susceptible triticale cultivar ‘Timbo’. Gene expression is displayed as log2-transformed TPM values. Values for avirulence genes are indicated as black dots. There are several gene copies of *AvrPm17* and *AvrPm3*^*a2/f2*^ in the THUN-12 genome. *AvrPm17* gene copies differ by two SNPs and are represented separately. Two of the *AvrPm3*^*a2/f2*^ copies are identical and represented (v1,v2) as a single data point, whereas the third copy carries a single synonymous SNP (v3) and is represented separately **B)** phenotypes of isolate THUN-12 at 10dpi on wheat differential lines or transgenic lines containing *Pm17* in the ‘Bobwhite’ background **C)** Result of the single interval QTL mapping approach, based on 55 randomly selected progeny of the mapping population Bgt_96224 X THUN-12 on the *Pm3c* containing NIL Sonora/8*CC. The black line indicates the logarithm of the odd score (LOD score) of the association. The dashed red line indicates the significance threshold estimated by 1000 permutations. **D)** represents the region corresponding to the 1.5LOD interval in the two genomes of isolates Bgt_96224 and THUN-12. Genes are indicated by colored boxes. Gene orientation is indicated by arrows. Members of different candidate effector families are represented by different colors as indicated.

In summary, we present a chromosome-scale genome assembly of the *B.g. triticale* hybrid pathogen based on significant advances in long-read sequencing technology and a high-resolution genetic map thereby allowing a molecular analysis of the hybrid genome structure of a fungal plant pathogen. The *B.g. triticale* genome is defined as a mosaic between two highly collinear parental genomes of wheat and rye powdery mildew and includes several lineage specific regions harbouring highly expressed candidate effector genes. The reference genome of *B.g. triticale* provides the basis for future identification and functional validation of virulence factors involved in host-adaptation on triticale. We furthermore provide evidence that avirulence and suppressor genes identified in wheat mildew are fully functional in *B.g. triticale* and thereby exemplify the importance of genomic analyses of plant pathogens for resistance breeding in triticale. We propose to pre-emptively combine genomic resources and pathogen diversity analyses to increase the efficiency of resistance breeding in triticale and beyond.

