## Supplementary material for "Host adaptation through hybridization: Genome analysis of triticale powdery mildew reveals unique combination of lineage-specific effectors": Methods_SupplementaryMaterial

### Material and Methods

#### Fungal isolates, mapping population, DNA extraction

The *B.g. triticale* isolate THUN-12 was described in (Menardo et al., 2016). Fungal isolates were maintained in the asexual state at 2°C on leaf segments of susceptible cultivar Kanzler and propagated to new leaf segments every 2 months. Crossing of parental isolates, genotyping and genetic map generation of Bgt_96224 X THUN12 was described in (Müller et al., 2019). High molecular DNA extraction for PacBio sequencing was performed according the protocol described in (Bourras et al., 2015)

#### Sequencing, Assembly, Polishing

PacBio sequencing was performed with PacBioSequel technology at the Functional Genomic Center Zurich (FCGZ) on three SMRT cells. The assembly was created with HGAP4 (SMRTLink Version 6.0.0.47841) (Chin et al., 2013) with default parameters (genome size was set to 166Mb based on the size estimation in (Müller et al., 2019) The assembly was subsequently polished using the Arrow software implemented in HGAP4. Prior to scaffolding an additional round of assembly polishing was performed based on Illumina sequencing of isolate THUN-12. To do so, Illumina reads trimmed using Trimmomatic (v 0.38, (Bolger et al., 2014), options; TruSeq3-PE-2.fa:2:30:10 LEADING:10 TRAILING:10 SLIDINGWINDOW:5:10 MINLEN:50 and subsequently mapped to the contigs of the THUN-12 assembly with bwa mem -M (v0.7.17, ), followed by the SAMtools commands fixmate, sort and markdup (v 1.7 (Li et al., 2009) and picards (vXX, https://github.com/broadinstitute/picard). The subsequent polishing was performed using pilon (v1.23, (Walker et al., 2014) with default parameter. The polished contigs were scaffolded into chromosomes using a genetic map. To do so, re-sequenced progeny of the biparental cross Bgt_96224 X THUN-12 were mapped on the THUN-12 contigs and production of the genetic map and subsequent scaffolding was performed as described in (Müller et al., 2019). Contig that contained mis-assemblies were split at the middle position between the two closest markers. To mark sequence gaps in the chromosome assemblies, stretches of 200 Ns were included. Lastly, chromosomes were oriented and numbered to match the chromosome of wheat powdery mildew isolate Bgt_96224 (Müller et al., 2019)

#### Annotation and effector definition

Annotation was performed using maker (v2.31.9, (Cantarel et al., 2008) based on the protein sequences of Bgt_96224 (Bgt_CDS_2018_39, Müller et al. 2019) and *B.g hordei* isolates DH14 and RACE1 (Frantzeskakis et al., 2018a) using the protein2genome=1 option. Repeat sequences were masked using a custom repeat library (nrTREP19, PTREP19, https://botserv2.uzh.ch/kelldata/trep-db/index.html) as well as with enabling the model_org=fungi option for the RepBase masking with the Repeatmodeler. In a parallel approach we identified the core Ascomycete genes with BUSCO 4.0.6, lineage asocmycota_odb10,

(Simao et al., 2015) in the DH14 proteins (Frantzeskakis et al., 2018b) and projected 132 missing genes that were annotated in DH14 but not in THUN-12 using maker onto the THUN12 genome assembly.

To define putative candidate effector genes we took advantage of the previously defined effector annotation (Müller et al., 2019). We aligned all proteins of THUN-12 against the protein sequence of the updated candidate effectors in Bgt_96224 (Bgt_CDS_v4_20) using blastp (v2.6.0+, (Camacho et al., 2009) (cutoff, bit>100) and assigned the gene to the corresponding candidate effector family. SignalP5.0 (Armenteros et al., 2019) was used to predict signal peptides for the remaining genes. Proteins containing a signal peptide were aligned against the proteins of *Podospora anserina* (https://www.ncbi.nlm.nih.gov/genome/?term=Podospora+anserina+S+mat%2B+genome, accessed 20.03.2020) and *Neurospora crassa* (https://www.ncbi.nlm.nih.gov/genome/?term=Neurospora+crassa+OR74A, accessed 20.03.2020) with blastp (v2.6.0+, (Camacho et al., 2009). Proteins with significant hit (cutoff, bit>100) were not considered as candidate effectors. The remaining proteins were aligned against the non-effectors gens of Bgt_96224 blastp (v2.6.0+, (Camacho et al., 2009) and the remaining 48 that had no significant hits to non-effector gene in Bgt_CDS_v4_20) were added to the list of candidate effectors in THUN-12. An additional two proteins had no significant hit to non-effector genes but did not contain a start codon and were therefore excluded. The list of candidate effectors can be found at https://github.com/MarionCMueller/B.g.-triticale-isolate-THUN-12/tree/master/Candidate_effectors.

**Transposable element annotation**

Transposable elements were annotated using the RepeatModeler v2.0 (http://www.repeatmasker.org/) software based on the non-redundant TREP nucleotide sequence database nrTREP19 database available at https://botserv2.uzh.ch/kelldata/trep-db/index.html. Based on the RepeatModeler output we used BEDtools complement to calculate the proportion of the THUN-12 genome that is free of repeats. The most abundant TE superfamilies according to the classification in (Wicker et al., 2007) were extracted and visualized with ggplot2 (Wickham, 2009) in R (Team, 2008).

**Determination of genomic origin**

The genomic origin of the chromosome of *B.g. triticale* THUN-12 was determined according to the rational described in (Menardo et al., 2016). Genomic resequencing data of *B.g. secalis* and *B.g. tritici* and THUN-12 ((Menardo et al., 2016) isolates were mapped against the THUN-12 genome as described in(Müller et al., 2019). Subsequently, variant calling was performed simultaneously on all isolates using freebayes (v1.1.0-54-g49413aa, https://github.com/freebayes/freebayes) with the freebayes-parallel script with the -p

1 -m 8 options. The resulting vcf file filtered with VCFtoosl for sites with quality above 20 and genotypes with a min coverage of 15. Only sites without missing genotype information were retained with the max-missing 1 option and the vcf file was transformed to genotypes using the --extract-FORMAT-info GT command (Danecek et al., 2011). The genotype file was subsequently analysed in R (Team, 2008). For the analysis only variants that are identical in all isolates within one f.sp and different in the other f.sp. were considered as fixed for this f.sp. and the origin of that site in *B.g. triticale* THUN-12 was assigned according to these informative sites. To estimate the contribution of each parental lineage to the THUN-12 genome, the recombination breakpoints between *B.g. tritici* and *B.g. secalis* segments were defined as the middle position between the two closest informative markers on the chromosome. The total contribution per chromosome was estimated by adding the segments originating from *B.g. secalis* and *B.g. tritici*, respectively.

**Analysis of genomic rearrangements**

Whole genome comparison was performed using NUCmer pipeline implemented in the MUMmer software (v4.0.0, (Marcais et al., 2018). Alignments were visualized using the mummerplot command with the --filter --color option. Plots were subsequently generated using gnuplot v5.3 (http://www.gnuplot.info/). To create an estimate of alignment breakpoints that are part of an inversion, duplication or gap we processed the NUCmer .delta output with the MUMmer show-diff program. The output of the show-diff program was subsequently parsed using a custom R script to extract putative rearrangements that are larger than 1kb and smaller than 10kb. For the detailed visualization of individual gene loci, the delta file was processed with dnadiff and the resulting .1coords file was used to extract alignment position.

For the comparative analysis of rearrangements located in regions inherited from *B.g. secalis* and *B.g. tritici* in THUN-12, only putative rearrangements larger than 1kb were considered. Locations of putative inversions, deletions and gaps were extracted from the mummer output form as described above. Rearrangements overlapping with segments inherited from *B.g.secalis* were identified using the BEDTools intersect (v2.26.00, (Quinlan and Hall, 2010) command. To test for enrichment of rearrangements inherited from *B.g. secalis*, we performed a $X^2$-goodness-of-fit test with the R base (v4.0.2, (Team, 2008)) function chisq.test(), p-values<0.05 were considered significant.

**Phenotyping**

Phenotyping was performed as described in (Bourras et al., 2019) on detached leaf segments of the following wheat differential lines: Axminster/8*CC (Pm1a), Ulka/8*CC (Pm2), Asosan/8*CC (Pm3a),

Michigan Amber/8*CC (Pm3f), Chul/8*CC (Pm3b), Sonora/8*CC (Pm3c), Kolibri (Pm3d) and the transgenic line Pm17#181 (Singh et al., 2018; Müller et al., 2021). Susceptible wheat cultivars Kanzler and Chancellor (CC) and appropriate Bobwhite sister lines were used as infection controls. For the QTL mapping the leaf coverage of progeny of the cross Bgt_96224 X THUN-12 was scored based on the following phenotypic scale virulent = 1, intermediate/virulent = 0.75, intermediate = 0.5, avirulent/intermediate = 0.25, avirulent = 0. For the final phenotypic score, values of at least 4 independent leaf segments were averaged.

**QTL mapping**

QTL mapping was performed as described in Müller et al. 2021 using the R package Rqtl (v1.46-2) (Broman et al., 2003). In brief, phenotyping data and map position of the cross Bgt_96224 X THUN-12 were imported using the read.cross() command. Genotyping data were processed using jittermap() and calc.genoprob() prior to QTL mapping with scanone(, model=”np”). Significance threshold was estimated with 1000 permutation using the scanone(model=”np”, n.perm=1000) command. The significance interval was extracted using the lodint(,extendtomarkers=TRUE) command.

**Annotation of avirulence gene loci**

To compare the avirulence loci between the two isolate Bgt_96224 and THUN-12, gene annotations and expression levels were visualized using the integrative genome browser IGV (v2.8.10). RNAseq-data of isolate Bgt_96224 on susceptible wheat cultivar ‘Chinese Spring’ and THUN-12 on susceptible triticale cultivar ‘Timbo’ were generated previously (Menardo et al., 2016; Praz et al., 2018). RNAseq-data was mapped to the respective genome using the method described in (Praz et al., 2018). For the *Avr* loci, annotation of low-quality genes that show no transcriptional support or exhibit homology to transposable elements were removed for better visualization. Information about genomic alignment was exported from the MUMmer ouput .1coords file (see above). The loci were visualized using with the ggplot2() (Wickham, 2009) package in R ((Team, 2008).

**Gene expression analysis**

Gene expression analysis of isolate THUN-12 was performed on RNAseq-data that was generated previously (Menardo et al., 2016) from infected triticale (cv. Timbo) at two days post-infection (2dpi). This time point corresponds to the establishment of the haustorial feeding structure of the fungus (Praz et al., 2018). Gene expression was quantified using salmon (v1.3.0) (Patro et al., 2017). The salmon index command was used to build the index from the CDS file. Subsequently, command salmon quant -l A -validateMapping was used to quantify the number of reads per transcript. TPM (transcript per million) values for each gene was extracted from the salmon output file and log2 transformation was performed in R (v4.0.2) with log2() command (Team, 2008).

**Identification of lineage specific genes**

Presence/absence polymorphisms of genes were estimated based on the method described in (Müller et al., 2019). In brief, genomic coverage per base was extracted from genomic mapping data using the SamTools (v1.7, (Li et al., 2009) depth -a command. Coverage per gene was obtained by averaging the values for each base for the positions between start and stop codon (including intron). Normalized values per gene were obtained by normalizing individual gene coverage by the average coverage of all genes. Gene with a normalized coverage value lower than 0.1 were considered absent from the isolate. Genes were only considered absent from one fungal lineage if the gene was absent in all analyzed isolates of this lineage. As an additional control, candidate effectors specific for the *B.g. secalis* lineage that were identified with this method, were further aligned to the Bgt_96224 genome (Bgt_genome_v3_16, (Müller et al., 2019) with blastn and plastp (BLAST+, v2.2.31+ (Camacho et al., 2009)) to verify their absence in the wheat mildew genome.

Lineage-specific effectors and structural variation in the underlying loci of the THUN-12 and Bgt_96224 genomes was visualized using the ggplot2() (Wickham, 2009) package in R (v4.0.2, (Team, 2008)), based on the following resources: Alignment information of the two genomes of THUN-12 and Bgt_96224 were extracted from the MUMmer output .1coords file (see above). Single copy orthologs were defined based on proteome data of Bgt_96224 (Bgt_CDS_v4_20) and THUN12 (THUN12_CDS_v1_2) using OrthoFinder (v2.4.1. (Emms and Kelly, 2019) with default parameters.

**Protein modelling**

For protein modelling, the signal peptide of selected candidate effectors were predicted with SignalP5.0 (Armenteros et al., 2019). Protein structure prediction was performed on the mature protein sequence (without signal peptide) with INTFOLD5.0 (McGuffin et al., 2019). Domain prediction for the best multi-template model was extracted, colored and visualized using JalView software (v2.11.1.3, (Waterhouse et al., 2009) The five best multi-templates models predicted by INTFOLD5.0 were extracted and deposited at GitHub repository https://github.com/MarionCMueller/B.g.-triticale-isolate-THUN-12/tree/master/Protein_modelling.

**Data availability**

Assembly and annotation of the THUN-12 isolate created in this study are available at ENA under accession number (PRJEB40658). PacBio raw reads are available at SRA under accession number PRJNA666512. Resequencing data of the mapping population Bgt_96224 X THUN-12 was generated previously (Müller et al., 2019) and is available at SRA (PRJNA453697). Assembly and annotation of Bgt_96224 (Bgt_CDS_v4_20) can be found at ENA under accession number PRJEB42580. RNAseq-data of isolate THUN-12 on susceptible triticale cultivar (cv. Timbo) was generated previously (Menardo et al., 2016) and can be found on SRA PRJNA427159. Re-sequenced isolates were published previously (Menardo et al., 2016)used in this study are available at the SRA under accession number PRJNA290428.

**Note S1 Candidate effector gene definition**

Definition of candidate effectors in plant pathogenic fungi remains challenging, mainly because the majority of candidate effector genes do not show homology to functionally characterized protein domains (Lo Presti et al., 2015). The absence of domain annotation was previously also described for the effector complement of *Blumeria graminis* (Menardo et al., 2017). A recent characterization of candidate gene effectors in the chromosome-scale assembly of wheat mildew isolate Bgt_96224 showed that candidate effector genes, apart from encoding a signal peptide, are typically part of effector families that are shared between the *B.g. tritici* and *B.g. hordei* lineage but are not present in other fungi such as the non-pathogenic species *Podosphera anserina* and *Neurospora crassa* (Müller et al., 2019). Because candidate effector families are ancient and present in several ancient lineages of *Blumeria graminis* we used a homology-based approach based on blast searches to identify genes in THUN-12 that show similarities to the candidate effector families of *B.g. tritici*. With this approach we identified 964 genes that show homology to members of the previously identified candidate effector families (Müller et al., 2019). A total of 953 genes in THUN-12 exhibited significant similarities with members of a single effector family and could therefore be unambiguously assigned. The remaining 11 genes had blast significant hits to members of several effector families, these genes were assigned to the effector family with the strongest hit. To account for the possibility that the THUN-12 annotation could contain candidate effectors that are not annotated in Bgt_96224 or candidate effectors that show no homology to effectors in Bgt_96224, we performed an additional round of candidate effector gene identification based on the remaining genes found in the THUN-12 genome, focusing on genes with a predicted signal peptide (SingalP5.0). We found an additional 47 genes that contained a signal peptide and were absent from the non-pathogenic *Podosphera anserina* and *Neurospora crassa* genomes. These genes were added to the list of candidate effector genes, resulting in a final list of 1,011 candidate effectors. The complete list of putative candidate effector genes in THUN-12 and their effector family affiliation is available on the *B.g. triticale* THUN-12 github repository:
https://github.com/MarionCMueller/B.g.-triticale-isolate-THUN-12/tree/master/Candidate_effectors

#### Note S2 Lineage-specific effector gene expansions

As part of this project on the wheat/rye powdery mildew hybrid *B.g. triticale,* we were confronted with the challenge that a high quality genome of the wheat powdery mildew parental lineage was available whereas there was no comparable resource for *B.g. secalis* (Müller et al., 2019). Therefore, an approach based on the two high quality genomes of triticale and wheat powdery mildew would only allow to detect putative lineage-specific effectors inherited from the rye lineage. To circumvent this shortcoming, we expanded our previously established coverage-based approach to predict the presence/absence of a particular candidate effector gene in THUN-12 based on the resequencing data of several isolate from both formae speciales

(Müller et al., 2019). To do so we calculated the normalized coverage per gene and considered genes with a normalized coverage below 0.1 as absent from a particular isolate. Only candidate effectors that were absent in all the analyzed isolates were considered absent from this particular lineage. Using this approach, we found 22 genes that were absent from the *B.g tritici* lineage, among those 9 candidate effector genes (Figures 1, S3). Vice-versa, 191 genes were absent from the *B.g. secalis* lineages, among those 109 candidate effectors (Figures 1, S4).

**Note S3 Detailed description of *B.g. secalis* lineage-specific effector clusters**

Using a coverage-based approach to detect lineage specific effectors in the *B.g. triticale* isolate THUN-12 we identified five genomic loci that harbor the nine *B.g. secalis* lineage specific effector genes (Figure 1). Among the five, the three loci located at the telomeres of Chr-01, Chr-4, Chr-11 were particularly interesting, whereas the loci on Chr-09 and Chr-08 contained a single lineage-specific effector that was expressed at low levels during haustorium formation (i.e. 48hpi) and might therefore be less relevant for the establishment of infection (Figure 2D,E, Figure S5A). The locus identified on Chr-01 contains 15 effectors from seven different candidate effector families in the ~800kb interval inherited from *B.g secalis,* whereas the corresponding region in *B.g tritici* isolate Bgt_96224 contains only 10 candidate effectors (Figure 2A). Using the MUMmer output and synteny analysis of the effectors and single copy orthologous genes in this locus we found evidence for an inversion that affects five genes, among those a candidate effector from family E030. The most striking difference between THUN-12 and Bgt_96224 in this locus was however that *BgTH12-04329*, the effector for which we detected the lineage-specific signal. This gene is present in four very similar copies in the THUN-12 genome (*BgTH12-04325/27/29/31),* whereas there is a single effector in Bgt_96224 (BgtE-5973) (Figure 2A,S5B,C). Interestingly a second candidate effector is also present in four copies (BgTH12-04324/26/28/30, family E001), whereas the corresponding region in the Bgt_96224 genomes contains a single copy of the gene (BgtE-20121). To confirm that this candidate cluster expansion is indeed inherited from *B.g. secalis*, we analyzed the coverage across the four copies (*BgTH12-04324/26/28/30* and *BgTH12-04325/27/29/31*) in the *B.g. tritici* and *B.g secalis* isolates (Figure S5B). The sum of coverage across the genes indicates that *B.g. tritici* indeed only has one gene, whereas the sequenced *B.g. secalis* isolates all have five or more copies of these effectors (Figure S5B). We therefore concluded that there is indeed a cluster expansion that was inherited form *B.g. secalis. BgTH12-04329* and its paralogues/orthologous encode for small (120 amino acids) proteins with a predicted signal peptide, several cysteine residues and is predicted to model to a RNAse-fold (Figure S5C,D). This RNase-fold was predicted for many *Blumeria* effectors including all the identified avirulence genes both in *B.g. tritici* and *B.g. hordei* (Bourras et al., 2019; Pennington et al., 2019; Hewitt et al., 2020; Bauer et al., 2021; Müller et al., 2021). In addition, *BgTH12-04329* and its orthologues are part of the top 5% of all expressed genes in

THUN-12 during haustorium formation (48hpi). High expression levels of effector genes during early infection timepoints has been described for most functionally characterized virulence and avirulence genes in Blumeria (Praz et al., 2017) and therefore suggests an important function during establishment of the infection (Figure S5A).

The locus on Chr-04 contains a cluster of genes belonging to effector family E029, with nine members in THUN-12 and six members in Bgt_96224 (Figure 2B). Again, we found evidence for several inversions in this locus. For instance, BgTH12-06220 is inverted compared to its likely orthologues BgtE-20041 in Bgt_96224 (Figure 2B). In addition, we found two identical candidate effector genes (BgTH12-06225/ 27) in THUN-12 (Figure S2B) that are absent in Bgt_96224 and all other *B.g. tritici* isolates. The two effectors BgTH12-06225 and BgTH12-06227 and y are among the highest expressed genes in THUN-12 (Figure S5A) and are predicted to contain a RNAase-fold (Figure S5E). In addition, we identified a lineage-specific gene BgTH12-06232 in THUN-12 that is not associated to any candidate effector family.

A particularly striking example of a genomic region containing lineage-specific effectors is located in the telomeric region on Chr-11. This segment is absent in wheat powdery mildew Bgt_96224. We found that THUN-12 encodes for seven candidate effectors, whereas Bgt_96224 encodes for only three effectors in this region. One candidate effector belonging to effector family E003 was present in THUN-12 but is not annotated in Bgt_96224 because it is pseudogenized (Figure 2C). Moreover, a part of this *B.g secalis* inherited segment was absent from the Bgt_96224 assembly (Figure S2,2C). To rule out that this region was not integrated in the Bgt_96224 assembly, we compared the segment with the unassembled fraction of the Bgt_96224 assembly but did not detect any matching region. Combined with the fact that our initial coverage-based approach did not detect any mapping reads to these genes in *B.g. tritici* isolates we therefore excluded the possibility of a scaffolding error in Bgt_96224 and concluded that this region indeed is specific to the *B.g. secalis* lineage. Interestingly the region contains three candidate effectors that all belong to family E003. Furthermore, we found that two of these effectors *BgTH12-06788/ BgTH12-06790* are again among the highest expressed gene in THUN-12 (Figure S5A) and are modelled into an RNAase-fold (Figure S5F).

**Note S4 Comparison of lineage-specific effectors to known candidate effector gene families**

The effector complement of *Blumeria graminis* has been characterized previously based on different genomic and transcriptomic resources (Pedersen et al., 2012; Menardo et al., 2017; Frantzeskakis et al., 2018b; Praz et al., 2018; Müller et al., 2019). It was established that candidate effectors exhibit increased levels of presence/absence polymorphisms, copy number variation, transcriptional plasticity and increased diversity compared to non-effector genes. Interestingly, the rate of these properties varied between different effector families (Pedersen et al., 2012; Müller et al., 2019) and was particularly pronounced in effector families that are small (100–200 aa) and highly expressed. These small and highly expressed candidate effector families were termed group 1 effector in (Müller et al., 2019). Most importantly, many of the virulence effector cloned in *B.g. hordei* as well as all cloned avirulence genes in *B.g. tritici* and *B.g hordei* are part of these so called group 1 effectors (Müller et al., 2019).

Many of the candidate effector genes that were identified in our search for lineage-specific effectors (Notes S3) fall into the category of group 1 effectors or show characteristics of group 1 effectors (Müller et al., 2019). For instance, family E003, that contains several lineage-specific effectors from *B.g. secalis* and *B.g. tritici*, is a large group 1 effector family that contains *AvrPm17* from *B.g. tritici (Müller et al., 2021)*, *Avra7* from *B.g. hordei* (Saur et al., 2019) as well as two described virulence effectors (BEC1016, CSEP0254) from *B.g. hordei* (Pliego et al., 2013; Ahmed et al., 2016). *BgTH12-05666* was found as lineage specific effector on Chr-08 and is part of family E004. E004 is a group 1 effector family and contains *AvrPm1a* form *B.g. tritici* (Hewitt et al., 2020) as well as a virulence effector from *B.g. hordei* (BEC1038) (Pliego et al., 2013). Family E029 contributes three *B.g. secalis* lineage-specific effectors, is located on the segment of Chr-04 and, was also classified as part of the group 1 effectors. None of the family members has been functionally characterized yet, but the gene *BgtE-20041* in Bgt_96224 is ranking second highest in a previously described screen for avirulence candidate effector genes, based on similarities to known avirulence genes from *B.g. tritici* and *B.g. hordei* (Bourras et al., 2019).

The family E135 was expanded in *B.g. triticale* and *B.g. secalis* (Figure 2A) and is represented by a single effector in Bgt_96224 (BgtE-5973) in the corresponding region on Chr-01 (Müller et al., 2019), Since the family contained a single family member in Bgt_96224 it could not be classified into either group 1 nor group 2 in Müller et al., 2019. However, the data presented in this study shows that this effector family contains more members in other *formae speciales*. The typical size of 125 aa as well the high expression of the effector genes furthermore suggests that E135 is also part of the group 1 effector gene families (Figure S5A,C).

In contrast to the above described group 1 candidate effectors, the five lineage-specific effectors belonging to family E001 (on Chr-01 and Chr-09, respectively), belong to the group 2 effectors. Group 2 effectors are generally larger (>300aa) and lower expressed on a family level than group 1 effectors (Müller et al., 2019). To our knowledge, none of the E001 family members from *B.g. tritici* or *B.g. hordei* have been characterized so far.

**Note S5 *B. g. tritici* specific lineages clusters co-localize with known avirulence gene loci**

Using the approaches described above (Note S3), we identified several regions that contain candidate effectors specific to the *B.g. tritici* lineage and are absent from the *B.g secalis* (Figure 1). Three of the regions that show the strongest signal overlap with the physical intervals of the previously identified *AvrPm1a* (Hewitt et al., 2020), *AvrPm17* (Müller et al., 2021) and *AvrPm*$^{a2/f2}$ (Bourras et al., 2015) effectors (Figure 1). We have recently characterized the *AvrPm17*-locus (Müller et al., 2021) and established that the effector gene cluster in this locus underwent several rounds of expansion in different *ff. ssp*. For instance, the E003 cluster containing *AvrPm17* expanded after the split from the *B.g hordei* lineage and predates the split of the *B.g. secalis/B.g. tritici* lineages. In the same locus, a cluster of 12 family E011 genes specifically expanded in *B.g. tritici* whereas it is represented by only 2 and 1 genes in the corresponding genomic region of *B.g. secalis* and *B.g. hordei,* respectively.

*AvrPm1a* is part of candidate effector family E004, and we have previously described that the *AvrPm1a* locus contains a cluster of 20 E004 family members on Chr-06. The coverage-based analysis again provided evidence that part of this cluster is expanded in the *B.g. tritici* lineage when compared to *B.g secalis*. It is however worth noting that the *AvrPm1a* effector itself is present in *B.g. secalis*. Sequence comparison to the avirulent AvrPm1a from B.g. triticale THUN-12 and functional consequences of putative amino acid polymorphisms of the *B.g. secalis* AvrPm1a on recognition by Pm1a will be subject to further studies. The third previously identified locus with strong signal of lineage-specific expansion in *B.g. tritici* is the *AvrPm3*$^{a2/f2}$ locus. In this case, the absence of the *AvrPm3*$^{a2/f2}$ gene from the *B.g. secalis* lineage has been previously described (Bourras et al., 2019). There are additional loci with strong signal for lineage-specific effectors, for instance, the short arm of chromosome 11 contains a large cluster of the E003 family that spans across the entire chromosome arms (Müller et al., 2019). We propose that a high-quality genome of the *B.g secalis* lineage would be highly valuable to resolve the question of effector expansion of these two formae speciales.

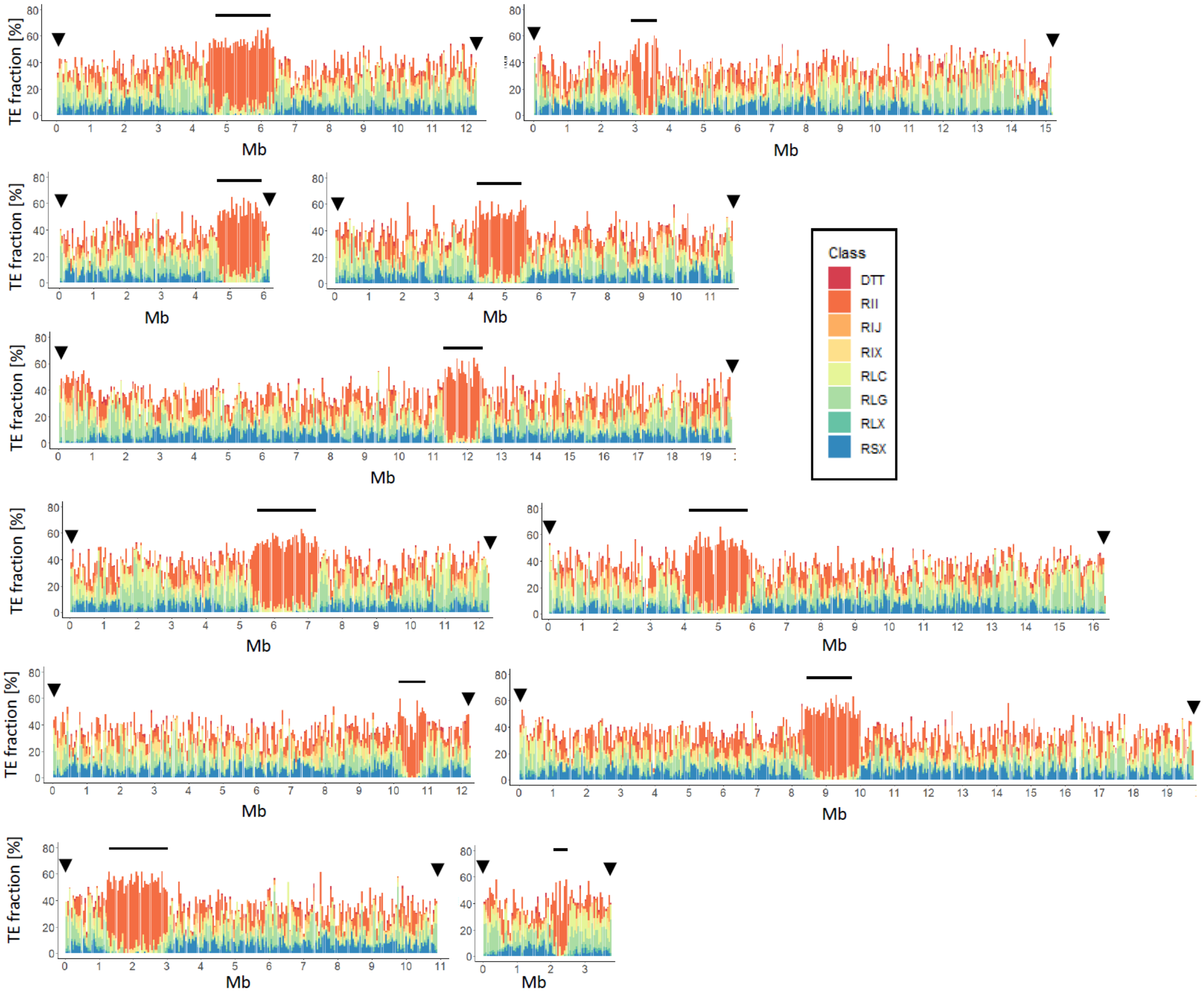


**Figure S1 Distribution of transposable elements along the eleven chromosomes of *B.g. triticale* THUN-12.** Transposable elements contribution was calculated in 50kb windows. The eight most abundant transposon superfamilies, classified according to the TE classification presented in (Wicker et al., 2007) are depicted. For each chromosome the genetic centromere is indicated with a black bar. The occurrence of the canonical telomeric repeat (TAACCC) is indicated by a black arrowhead.

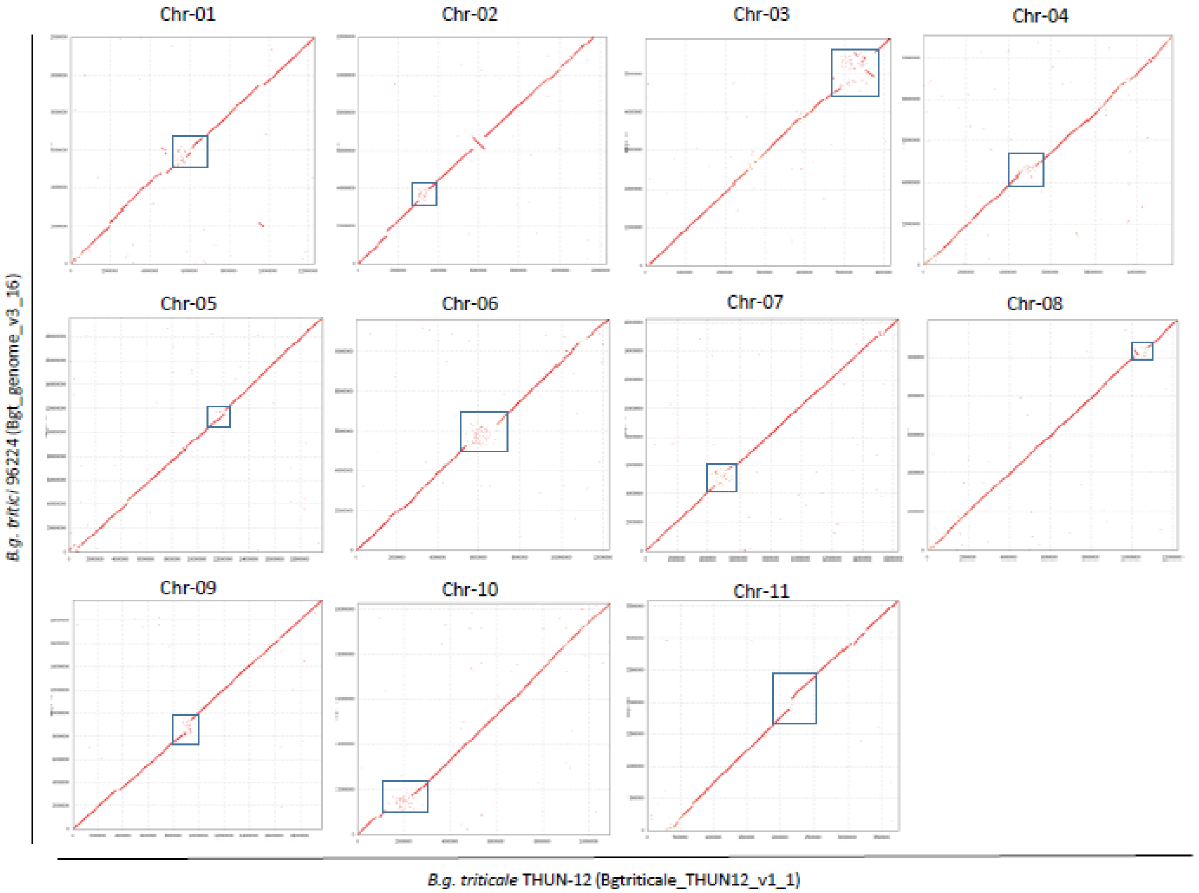


**Figure S1 Chromosomal alignment of the eleven chromosomes between *B.g. triticale* THUN-12 and *B.g. tritici 96224.*** Data was obtained from whole genome comparison with the MUMmer software. For each chromosome, the position of the centromere is indicated by a blue box.

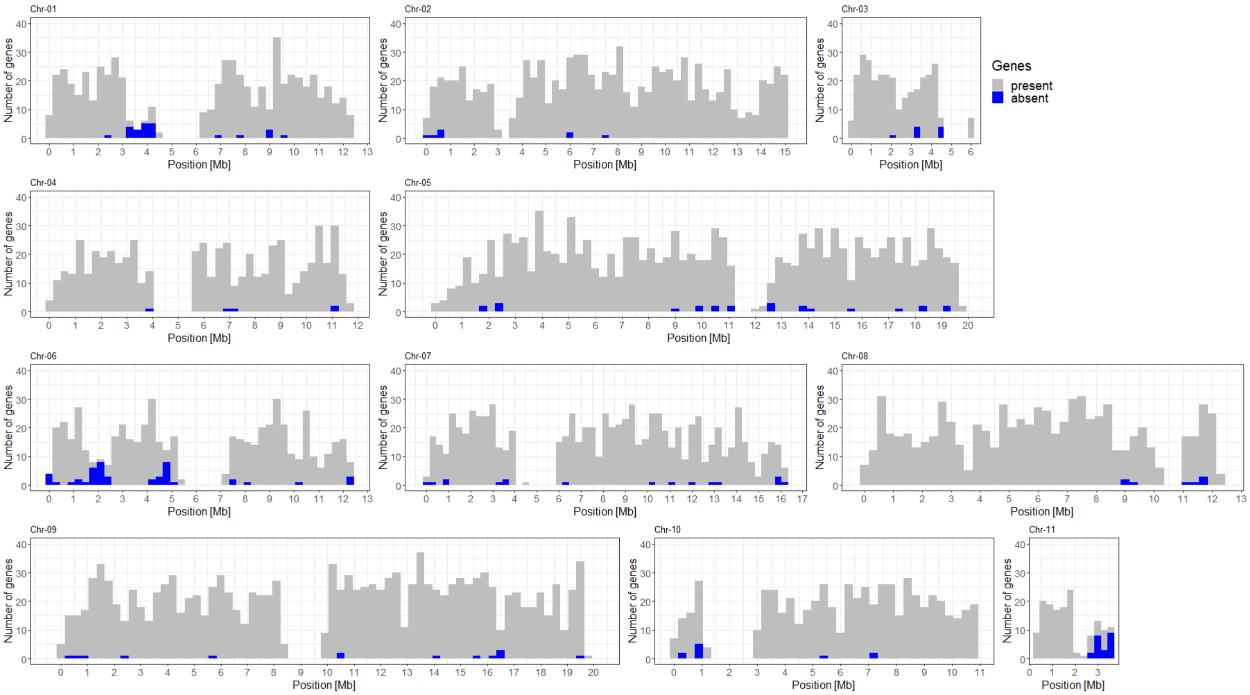


**Figure S2 Lineage-specific absences of genes in *B.g. secalis*.** Presence and absence of genes in the THUN-12 assembly in five *B.g. secalis* isolates were estimated based on genomic coverage of the genes. Genes were categorized as absent in the *B.g. secalis* lineage, if the gene was absent in all five isolates. Plots show gene density for each chromosome in 300kb non-overlapping windows. The presence and absence of genes in *B.g. secalis* is indicated as follows: grey indicates genes present in at least one of the five isolates, blue represents genes absent all five isolates.

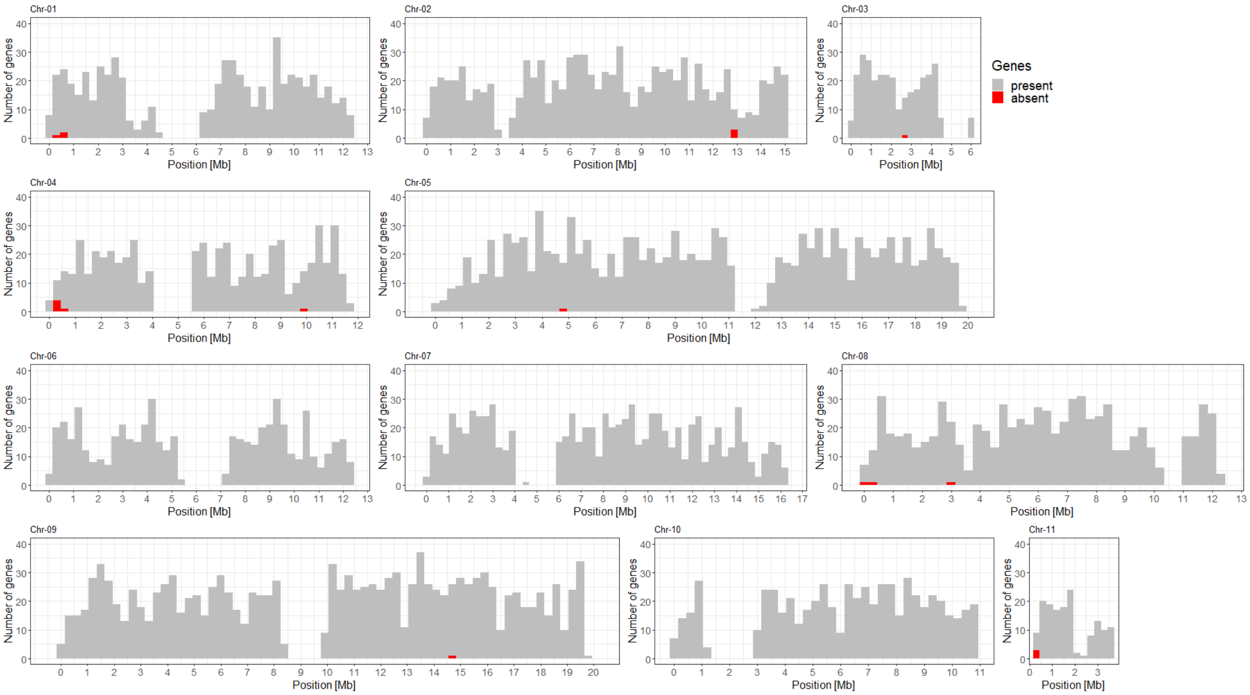


**Figure S3 Lineage-specific absences of genes in *B.g. tritici*.** Presence and absence of genes in the THUN-12 assembly in twelve *B.g. tritici* isolates were estimated based on genomic coverage of the genes. Genes were categorized as absent in the *B.g. tritici* lineage if the gene was absent in all twelve isolates. Plots show gene density for each chromosome in 300kb non-overlapping windows. The presence and absence of genes in *B.g. tritici* is indicated as follows: grey indicates genes present in at least one of the 13 isolates, red represents genes absent all 13 isolates.

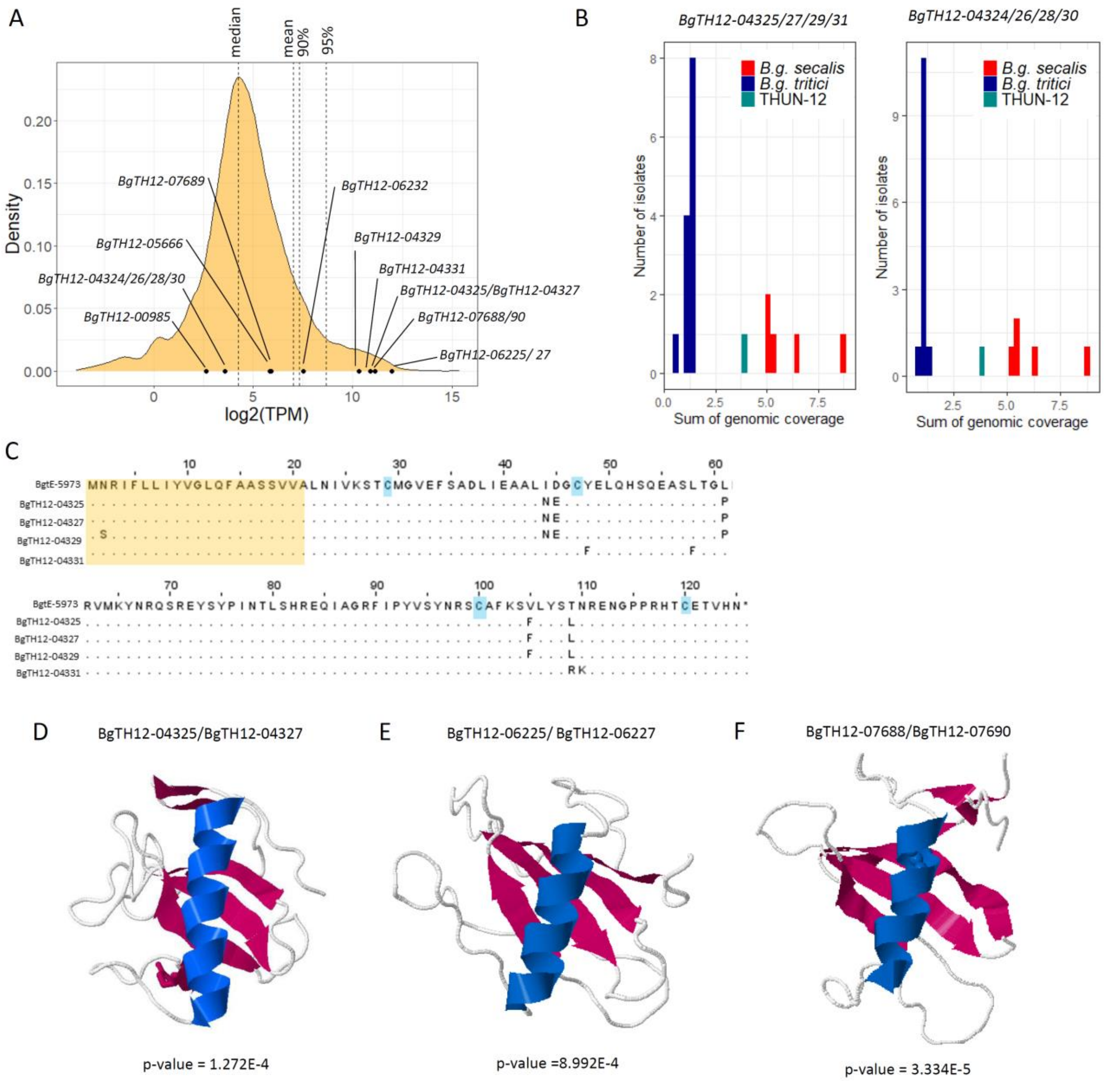


**Figure S4 Characterization of the lineage-specific effector genes inherited from *B.g. secalis* in the *B.g. triticale* hybrid THUN-12. A)** Expression of all genes in *B.g. triticale* THUN-12 are represented as log2(TPM) values. Expression levels of the lineage-specific effectors depicted in Figure 2 are indicated by a black dot. Identical genes are represented as a single value. **B)** Sum of normalized genomic coverage of the four genes belonging to candidate effector family E135 (BgTH12-04325/27/29/31) and E001 (BgTH12-04324/26/28/30) encoded in the effector cluster on Chr-01 **C)** Protein alignment of the E135 family members of Bgt_96224 and *B.g. triticale* THUN-12: The predicted signal peptide (SignalP5.0) is indicated by a yellow box. The four cysteine residues are indicated by a blue box. **D-E)** protein modelling of lineage specific effector genes using INTFOLD5.0. For each protein the predicted structure of the best model is depicted. Predicted α-helices were colored in blue, β-strands were colored in red. P-values of the best model is indicated below the figure for each model **D)** BgTH12-04325/BgTH12-04327 belongs to family E135. Significant model templates identified for this protein were the fungal ribonuclease: 1fusA, 1i0vA, 1rdsA, 3wr2A **E)** BgTH12-06225/BgTH12-06227 is part of family E029 and models to the following templates fungal ribonucleases: 1fusA, 1rmsA, 1futA and a putative protein kinase from *Candida albicans* 4c0tA , **F)** Candidate effectors BgTH12-07688/BgTH12-07689 belong to family E003, the family that contains *B.g. tritici* AvrPm17 and Avra7 from *B.g. hordei*. *BgTH12-07688/BgTH12-07690* models to the following templates: 6fmbA, the crystallized RNAase-like effector BEC1054 from *B.g. hordei* (Pennington et al., 2019)as well as fungal RNAases: 3whoA, 1fusA, 5gy6A.

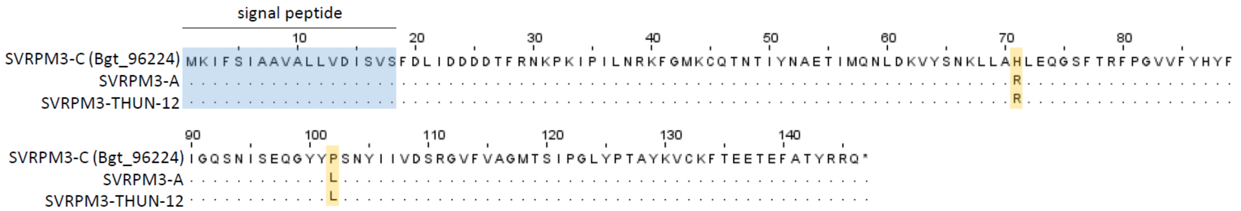


**Figure S5 Protein alignment of active and inactive SVRPM3 haplovariants from different isolates.** SVRPM3-C designated the previously described inactive SVRPM3 haplovariant encoded by isolate Bgt_96224, whereas SVRPM3-A is the active variant that is able to suppress Pm3-mediated resistance in wheat and was isolated previously in Pm3-virulent isolate Bgt_94202 (Bourras et al., 2015; Parlange et al., 2015). Isolate THUN-12 encodes for SVRPM3-A. The predicted signal peptide is indicated by a blue box. Polymorphic amino acid residues are highlighted by a yellow box.

#### Figures and Tables

Table S1 Summary and statistics of the scaffolded *B.g. triticale* THUN-12 assembly

| **Chromosome** | **Size** | **Gaps**[a] | **Sites *B.g. tritici***[b] | **BP *B.g. tritici***[c] | **% *B.g. tritici***[d] | **Sites *B.g. secalis***[b] | **BP *B.g. secalis***[c] | **% *B.g. secalis***[d] |
|---|---|---|---|---|---|---|---|---|
| TH12_chr-01 | 12’297’139 | 3 | 12’532 | 11’266’638 | 91.6% | 1’331 | 1’030’501 | 8.4% |
| TH12_chr-02 | 15’157’869 | 2 | 14’078 | 10’588’940 | 69.9% | 4’259 | 4’568’930 | 30.1% |
| TH12_chr-03 | 6’146’397 | - | 3’825 | 3’733’204 | 60.7% | 2’143 | 2’413’193 | 39.3% |
| TH12_chr-04 | 11’660’615 | 1 | 10’889 | 6’887’144 | 59.1% | 4’775 | 4’773’472 | 40.9% |
| TH12_chr-05 | 19’719’700 | 3 | 22’561 | 18’124’209 | 91.9% | 1’972 | 1’595’491 | 8.1% |
| TH12_chr-06 | 12’285’964 | - | 11’102 | 11’533’832 | 93.9% | 319 | 752’132 | 6.12% |
| TH12_chr-07 | 16’321’024 | 2 | 21’065 | 13’743’078 | 84.2% | 1’098 | 2’577’946 | 15.8% |
| TH12_chr-08 | 12’231’121 | 3 | 10’174 | 7’630’878 | 62.1% | 4’393 | 4’600’243 | 37.6% |
| TH12_chr-09 | 19’725’520 | 2 | 26’159 | 18’547’974 | 93.5% | 1’384 | 1’177’546 | 6.0% |
| TH12_chr-10 | 10’894’025 | - | 9’969 | 7’498’904 | 68.8% | 4’056 | 3’395’120 | 31.2% |
| TH12_chr-11 | 3’749’771 | 1 | 3’467 | 2’960’228 | 78.9% | 649 | 789’543 | 21.1% |
| Unanchored | 1’218’821 | - | - | - | - | - | - | - |
| Total[e] | 141’407’966 | 18 | 145’821 | 112’846’579 | 80.3% | 26’379 | 27’674’117 | 19.7% |

[a] Scaffolding gaps per chromosome, sequence gaps were marked by inserting a stretch of 200N into the sequence

[b] Number of fixed sites in THUN-12 that could be attributed to the parental f.sp.

[c] estimated genome proportion (in bp) contributed by each parental f.sp.

[d] estimated percentage of the chromosome length contributed by each f.sp.

[e] for the estimation of parental contribution the unanchored sequence was not considered

Table S2 Summary of rearrangements between the chromosome-scale assemblies of *B.g. triticale* THUN-12 and *B.g. tritici* Bgt_96224.

| **Type** | ***B.g. secalis*** | ***B.g. tritici*** | **Total** | **$X^2$p-values[a]** |
|---|---|---|---|---|
| Gaps | 28 | 71 | 99 | 1.86E-02 |
| Duplications | 1695 | 3063 | 4758 | 7.98E-188 |
| Inversions | 44 | 55 | 99 | 1.09E-10 |

[a] p-value of $X^2$-goodness of fit tests, p<0.05 indicates statistically significant deviation from the expected ratio given by the genomic proportion and indicates significantly more rearrangement in segments inherited from *B.g. secalis.*
